# Biaxial testing and sensory texture evaluation of plant-based and animal deli meat

**DOI:** 10.1101/2025.02.19.639170

**Authors:** Skyler R. St. Pierre, Lauren Somersille Sibley, Steven Tran, Vy Tran, Ethan C. Darwin, Ellen Kuhl

## Abstract

Animal agriculture is one of the largest contributors to global carbon emissions. Plant-based meats offer a sustainable alternative to animal meat; yet, people are reluctant to switch their diets and spending habits, in large due to the taste and texture of plant-based meats. Deli meat is a convenient form of protein commonly used in sandwiches, yet little is known about its material or sensory properties. Here we performed biaxial testing with multiple different stretch ratios of four plant-based and four animal deli meats and fit accurate material models to the resulting stress-stretch data. Strikingly, the plant-based products, turkey, ham, deli, and prosciutto, with stiffnesses of 378 ± 15 kPa, 343 ± 62 kPa, 213 ± 25 kPa, and 113 ± 56 kPa, were more than twice as stiff as their animal counterparts, turkey, chicken, ham, and prosciutto, with 134 ± 46 kPa, 117 ± 17 kPa, 117 ± 21 kPa, and 49 ± 21 kPa. In a complementary sensory texture survey, n = 18 participants were able to correlate the physical stiffness with the sensory brittleness, with Spearman’s correlation coefficient of *ρ* = 0.857 and *p* = 0.011, but not with the sensory softness or hardness. Notably, the participants perceived all four plant-based products as less fibrous, less moist, and less meaty than the four animal products. Our study confirms the common belief that plant-based products struggle to meet the physical and sensory signature of animal deli meats. We anticipate that integrating rigorous mechanical testing, physics-based modeling, and sensory texture surveys could shape the path towards designing delicious, nutritious, and environmentally friendly meats that mimic the texture and mouthfeel of animal products and are healthy for people and for the planet. Data and code are freely available at https://github.com/LivingMatterLab/CANN.

## 1. Introduction

Transitioning to a plant-based diet positively impacts both global and personal health [28, 37]. Consumer surveys have found that people tend to associate animal meat with *“delicious”* and plant-based meat with *“disgusting”* [21]. Matching the sensory experience of animal meat with plant-based ingredients is a major obstacle in convincing people to change their grocery shopping habits [6, 10, 21].

Animal deli meats are understudied for both mechanical and sensory textural properties [1, 16]. The primary research on deli meat is associated with the risk of listeria contamination, the presence of the bacteria in food, which is comparable to that of soft cheeses and packaged salads [3]. Animal deli meat is created through a restructuring process where multiple muscles are combined with chilled brine then formed to create a cylindrical shape, which is then easily cut into uniform, thin, round slices [23]. The resulting texture can vary significantly, depending on the processing methods and equipment, the addition of other ingredients including water, and the quality of the raw meat [16]. However, very little research has been conducted to analyze the textural differences of deli meat, likely because deli is a low cost product [1].

Although there are dozens of potential plant proteins that could be used to make plant-based meat analogs, soy, wheat, and pea protein are the most common [30, 36]. Plant-based meats designed to mimic minced or comminuted meat products are created commercially via extrusion or shear cell technology [17]. To date, no studies have simultaneously analyzed the *mechanical* and *sensory* properties of plant-based deli meats or provided a direct comparison between plant-based and animal deli meats.

The mechanical properties of meats and meat analogs are commonly evaluated by *texture profile analysis*, a double compression test that is supposed to mimic chewing two bites [29]. However, texture profile analysis works best with threedimensional samples with dimensions of one centimeter or larger [5], making it infeasible to evaluate the mechanical properties of a single prepackaged deli slice. To work around this limitation, a previous study used uncut deli poultry meats and then cut a one-centimeter thick slice for texture profile analysis [16]. This study also used the Warner-Bratzler and Kramer shear cell tests, which involve either one or multiple blades slicing through thick slices of deli meat [16]. Other options include mechanical tests using a rheometer to measure shear, but these are most effective for three dimensional materials like hotdogs and chicken [31, 29]. Uniaxial tension tests are only able to evaluate mechanical properties along a single material axis, so they are best suited for homogeneous materials like a tofurky deli slice [31]. To date, no deli meat study has used the classical mechanical test of *biaxial extension*, which is designed to measure the mechanical properties of thin sheets or membranes [12, 13, 20, 34], ideally suited to test single-slice deli meats.

The sensory texture properties of meat refer to the subjective perception of the mechanical properties experienced by human senses. Sensory texture properties can be evaluated by *expert sensory panels* or *consumer studies* [32], and properties includes primary parameters like hardness, viscosity, and springiness, and secondary parameters like chewiness and brittleness [32]. Although these mechanical characteristics may seem straight-forward, matching our sensory texture experience with mechanical testing remains challenging [32, 16]. For instance, while people can judge between hard, firm, and soft, an instrument has no sense of the boundaries between these categories [32]. Expert sensory panelists are given explicit instructions on what to evaluate with each bite of food, they usually have significant experience judging descriptive sensory properties, and they generally go through multiple rounds and hours of training sessions for each study [16, 4]. As a result of these strict requirements, most sensory panels have under twelve participants [4]. In contrast, consumer studies typically have over one hundred participants and do not require any particular training [7]. Consumer studies provide generalizable information on what sensory properties are liked or disliked, while sensory panels are thought to provide more specific, accurate, and actionable information to guide product development [7]. However, untrained consumers are still able to distinguish sensory texture properties, but with less precision than the trained panelists [2, 26].

In this work, we test four plant-based and four animal deli meats using biaxial extension to discover physics-based models that best characterize their behavior and identify their mechanical properties. To complement the mechanical characterization, we use untrained participants to evaluate which meats have noticeably different textural properties for the average consumer. Finally, we apply Spearman’s rank correlation between the mechanically derived stiffness and the textural sensory properties to understand how the quantitative and qualitative properties are correlated.

## 2. Methods

### 2.1. Mechanical testing

We test four plant-based deli meat products: oven roasted (Tofurky, Hood River, OR), hickory smoked (Tofurky, Hood River, OR), smoked ham style (Tofurky, Hood River, OR), and prosciutto style (Mia, Lakewood, NJ). For comparison, we also test four animal meat deli products: turkey breast (Hillshire Farm, Chicago, IL), chicken breast (Hillshire Farm, Chicago, IL), black forest ham (Hillshire Farm, Chicago, IL), and prosciutto (Columbus, Hayward, CA). Table 1 summarizes the ingredients of all eight products. For each meat type, we measure the sample thickness and test *n* = 8 samples in biaxial tension. Figure 1 shows our mechanical testing set-up, with rakes embedded in square samples of each deli meat.

**Table 1:**
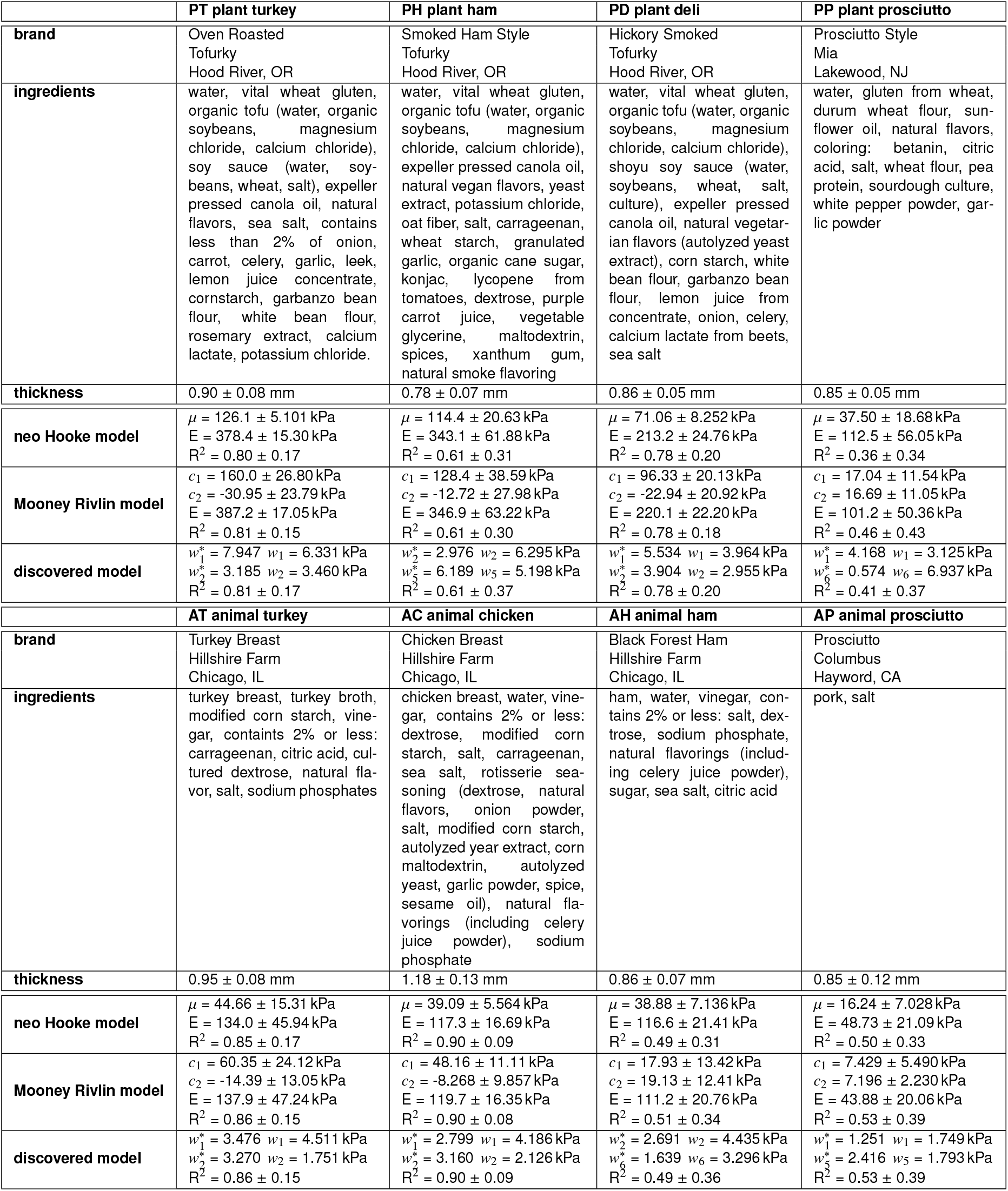
Plant-based and animal deli meat products. Products; brands; ingredients; thickness; neo Hooke parameter, stiffness, and goodness of fit R^2^; Mooney Rivlin parameters, stiffness, and R^2^; constitutive neural network parameters and R^2^. neo Hooke and Moony Rivlin parameters and stiffness are reported as the mean ± standard deviation of fitting each sample individually to the given model. Discovered model weights result from fitting the mean data from Table 1 and correspond to Figure 7 and equation (17). R^2^ is reported as the mean ± standard deviation of fitting the mean data from Table 1 across all five loading modes for each model.

**Figure 1:**
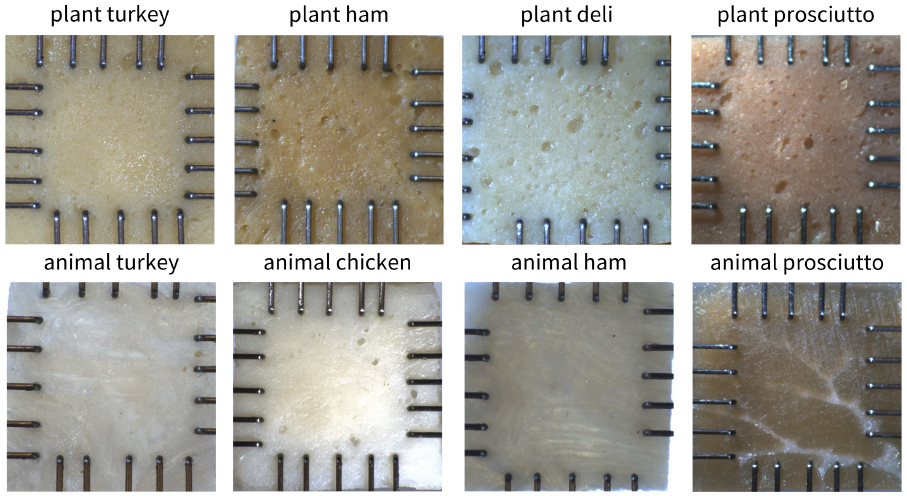
Samples of all eight plant-based and animal deli meats mounted for biaxial tension testing. The top row shows the plant-based meats, and the bottom row shows the animal meats. Rakes are inserted into the samples to form a 12×12 mm^2^ square shape. All plant-based meats display a homogeneous, isotropic microstructure with bubbles of different sizes randomly distributed across the samples, but with no apparent fibers. All animal meats display clearly visible fibers. In all animal turkey, chicken, and ham, fibers are not uniformly oriented in a preferred direction due to the manufacturing process, which combines multiple pieces of meat. In animal prosciutto, although fibers are visible, there is also no unique preferred fiber direction due to the random branching patterns of the fat throughout the muscle. We load all samples with random orientations.

### 2.2. Sample preparation and testing

We cut individual deli slices to approximately 15×15 mm^2^ sized samples and placed the sheets into the biaxial tester, the CellScale BioTester 5000 (CellScale, Waterloo, Ontario, CA). We gently push the 4×5 rakes into each sample [9], such that the rakes form a square grid of approximately 12×12 mm^2^, leaving an adequate sample overhang on each side, as shown in Figure 1. Then, we apply a small pre-load of 30 mN in both the x-and y-directions [18]. Pre-load is the amount of force required to stretch the sample to remove any slack, based on visual inspection of the force-displacement curve and the sample itself [35]. After applying the pre-load, we increase the biaxial tensile stretch quasi-statically at a rate of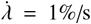. For all deli meats, except the plant-based and animal prosciuttos, we use the displacement-controlled setting to prescribe a maximum stretch of 10%. For the two prosciutto-type meats, we prescribe a maximum stretch of 25%, as these meats can sustain more stretch before exhibiting any signs of failure. For each sample, we perform five modes of tests in the same order with three cycles of stretch and recovery per test: strip-y, off-y, equi-biaxial, off-x, strip-x.

### 2.3. Stress and strain analysis

We process the data from the CellScale BioTester 5000 to obtain the average Piola stress for given stretch values *λ*_1_ and *λ*_2_ from the third stretch-recovery cycle of each of the five testing modes. First, we convert the force and displacement measurements to stresses and stretches. To do so, we measure the sample thickness *t* with calipers and set the gauge lengths *L*_1_ and *L*_2_ to the spacing between the tines in the 1- and 2-directions from the first step following the pre-load. Then, we compute the stretches *λ*_1_ and *λ*_2_ and the Piola stresses *P*_11_ and *P*_22_,

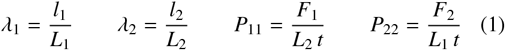

where *l*_1_ and *l*_2_ are the measured gauge lengths and *F*_1_ and *F*_2_ are the measured forces in the 1- and 2-directions. This results in five loading and five unloading curves for each of the ten samples. We resample and average all curves at equidistant stretch intervals to obtain an averaged stress pair {*P*_11_, *P*_22_} for each stretch pair {*λ*_1_, *λ*_2_}.

### 2.4. Kinematics

We characterize the deformation through the mapping ***x*** = φ(***X***) that maps a point ***X*** in the reference configuration to a point ***x*** the deformed configuration. We then describe the local deformation using the deformation gradient,

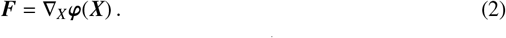

Multiplying ***F*** with its transpose ***F***^t^ introduces the symmetric right Cauchy Green deformation tensor ***C***,

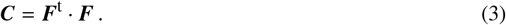

To characterize the deformation, we introduce the three invariants, *I*_1_, *I*_2_, *I*_3_,

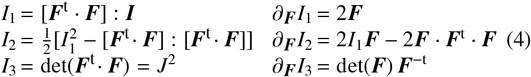

where ***I*** denotes the identity tensor. Since we are unable to measure the deformation in the thickness direction of the deli meats, we assume that the meats are perfectly incompressible, *I*_3_ = 1. In our biaxial extension tests, we stretch the sample in two orthogonal directions, *λ*_1_≥1 and *λ*_2_≥1. The incompressibility condition, 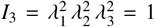, defines the stretch in the thickness direction as *λ*_3_ = (*λ*_1_ *λ*_2_)^−1^≤1. We assume that the deformation remains homogeneous and shear free, and the deformation gradient ***F*** remains diagonal,

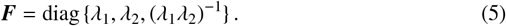

We can then specify the invariants (4) in terms of the biaxial stretches *λ*_1_ and *λ*_2_,

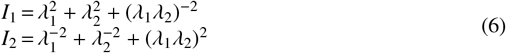

and explicitly take their derivatives,

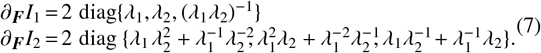

We assume a hyperelastic constitutive model, for which the stress, in our case the Piola stress ***P***, only depends on the current deformation state, in our case the deformation gradient ***F***, such that ***P*** = ***P***(***F***). To satisfy thermodynamic consistency, we can express the stress as a function of the strain energy density *ψ* as ***P*** = *≤ψ*(***F***)*/≤****F***, which we reformulate in terms of our set of invariants *ψ*(*I*_1_, *I*_2_, *I*_3_),

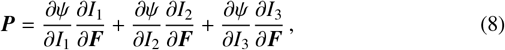

We explicitly enforce incompressibility by selecting the term in the third invariant as *ψ*(*I*_3_) = −*p* [*J*−1], such that *∂ψ / ∂I*_3_ *∂I*_3_*/∂****F*** = −*p* ***F***^−t^. Here *p* acts as a Lagrange multiplier that we determine from the zero-thickness-stress condition.

### 2.5. Biaxial testing

In biaxial extension tests, we stretch the sample in two orthogonal directions, *λ*_1_ ≥ 1 and *λ*_2_ ≥ 1, and, by incompressibility, *λ*_3_ = (*λ*_1_ *λ*_2_)^−1^ ≤ 1. We assume that the deformation remains homogeneous and shear free, and that the resulting Piola stress ***P*** remains diagonal,

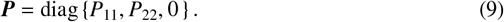

We use the isotropic first and second invariants 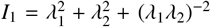 and 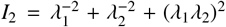 from equation (6) and their derivatives from equation (7) to determine the pressure *p* from the zero-thickness-stress condition in the third direction,

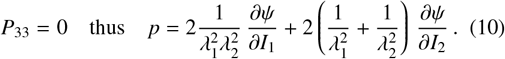

Equations (8) and (10) then provide an explicit analytical expression for the nominal stresses *P*_11_ and *P*_22_ in terms of the stretches *λ*_1_ and *λ*_2_ and the biaxial Piola stresses simplify to

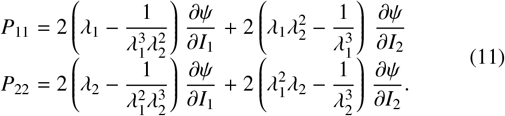

### 2.6. Mooney Rivlin and neo Hooke models

The Mooney Rivlin model is a two-term model with two parameters, *c*_1_ and *c*_2_,

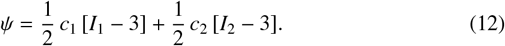

The explicit form of the Piola stresses (11) then simplifies to

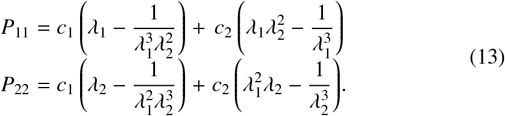

The neo Hooke model is a special case of the Mooney Rivlin model (12), where the parameter *c*_2_ is zero, leaving only a single term with one unknown parameter, *c*_1_,

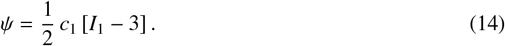

resulting in a simple expression for the Piola stresses (13) as

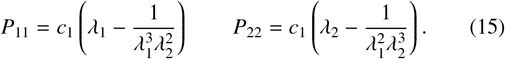

To determine the optimal parameter values, we minimize the following loss function using the least squares method,

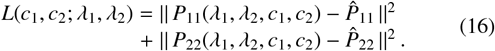

The Python script to optimize the Mooney Rivlin and neo Hooke parameters is included in the GitHub repository.

### 2.7. Automated model discovery

To confirm that the neo Hooke model or Mooney Rivlin model are the best models to describe the constitutive behavior of deli meat, or discover better suited constitutive models, we perform automated model discovery [14]. We use our custom-designed constitutive neural network that approximates a strain energy function in terms of the two invariants *I*_1_ and *I*_2_, and has16 parameters or network weights, 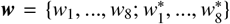, neight internal weights 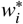 between its two hidden layers and eight external weights *w*_*i*_ out of its final hidden layer. We assume that the individual contributions to the free energy are fully decoupled. The free energy function for the two-fiber network takes the following explicit form [15],

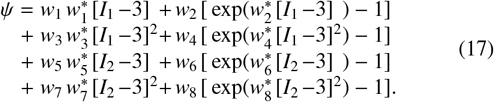

### 2.8. Model training

To discover models and parameters 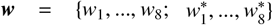 that best describe each meat product, we use the Adam optimizer to perform gradient descent on a weighted least squared error loss function *L* that penalizes the error between the discovered model ***P***(***F***_*i*_, ***w***) and the experimental data 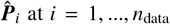 at *i* = 1, …, *n*_data_ discrete points, supplemented by *L*_*p*_ regularization [19],

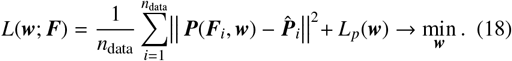

For the *L*_1_ regularization, we supplement the loss function by an *α*-weighted regularization term, 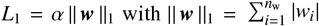.

### 2.9. Sensory texture survey

We prepare bite-sized samples of the eight deli meats, four plant-based, plant turkey, plant ham, plant deli, plant prosciutto, and four animal-based, animal turkey, animal chicken, animal ham, and animal prosciutto, as shown in Figure 3. No condiments or sauces are added to the deli slices.

We recruit *n* = 18 participants to participate in three surveys: the ten-question Food Neophobia Survey [25], the sixteen-question Meat Attachment Questionnaire [8], and our own twelve-feature Sensory Texture Survey. We also collect demographic information: age range, ethnicity, and gender. We instruct each participant to eat a sample of each meat product and rank its texture features according to our survey. The Sensory Texture Survey uses a 5-point Likert scale with twelve questions. Each question starts with “This food is …”, followed by one of the following features [22, 32]: *soft, hard, brittle, chewy, gummy, viscous, springy, sticky, fibrous, fatty, moist*, and *meaty*. The scale ranges from 1 for strongly disagree to 5 for strongly agree. Prior to beginning the survey, participants were reminded that viscosity is a fluid’s resistance to movement and the scale ranges from low viscosity, water-like, to high viscosity, peanut butter-like. This research was reviewed and approved by the Institutional Review Board at Stanford University under the protocol IRB-75418.

### 2.10. Statistical analyses

We use the correlation of determine, *R*^2^, to quantify the goodness of fit of the neo Hooke model, the Mooney Rivlin model, and the newly discovered constitutive models to the experimental biaxial data for each deli meat. *R*^2^ ranges from 0 for no correlation to 1 for a perfect correlation between model and experimental data. We use a one-way Analysis of Variance (ANOVA) with *p <* 0.05 to determine if the differences between meats for each of the twelve texture features are significant. To determine if the experimentally measured stiffness correlates with any of the texture features which are significant from the ANOVA, we use Spearman’s rank correlation. We report both the correlation coefficient *ρ* and the p-value for all significant correlations with *p <* 0.05.

## 3. Results

Table 1 provides a comprehensive overview of the eight products, their brands and ingredients, their thickness, and their constitutive model parameter values and fits. Of the four plantbased deli meats, plant turkey, plant ham, and plant deli are all Tofurky brand with the same first three ingredients: water, vital wheat gluten, and organic tofu. All four plant-based deli meats use wheat gluten as the primary plant protein. Of all eight meats, animal prosciutto has the smallest number of ingredients, with only pork and salt. As measured prior to loading the samples for biaxial testing, the thickness ranged on average from 0.78 mm for plant ham to 1.18 mm for animal chicken. Table 2 reports the mean of the loading and unloading curves across all *n* = 8 samples for each biaxial loading mode. The best fit parameters for the neo Hooke and Mooney Rivlin models are reported with the corresponding goodness of fit, *R*^2^. The stiffnes, *E*, is also reported for both models. All automatically discovered models have exactly two terms as a result of L1 regularization, with the reported weights corresponding to Figure 2 and equation (17).

**Table 2:**
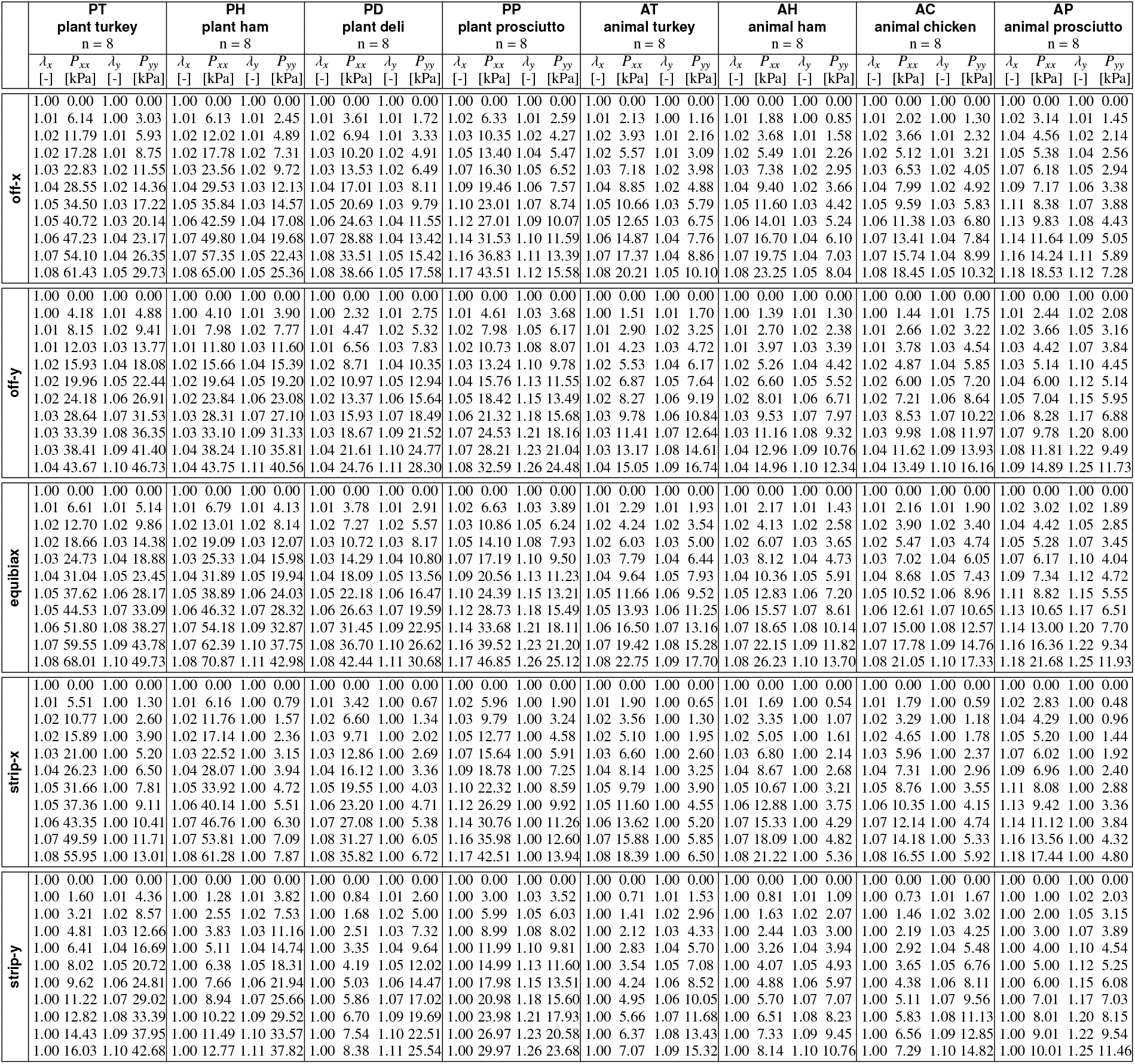
Biaxial testing data for plant-based and animal deli meat products. Mean stretch and stress for all five loading modes, abbreviated to eleven data points per loading mode. Stresses are reported as mean of loading and unloading curves for *n* samples for each biaxial loading mode: off-x, off-y, equibiax, strip-x, and strip-y. Extended data are available in the GitHub repository for this paper.

**Figure 2:**
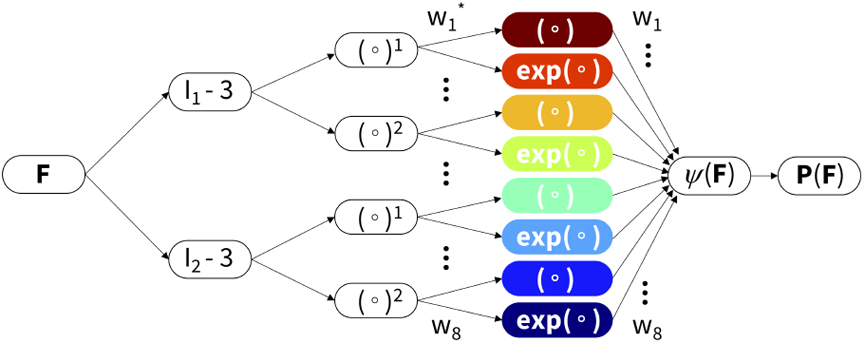
Automated model discovery. The model discovery uses an isotropic, perfectly incompressible constitutive neural network with two hidden layers and eight terms. The network takes the deformation gradient ***F*** as input and calculates its first and second invariant terms, [*I*_1_−3] and [*I*_2_−3]. The first layer generates powers of these invariants, (○)^1^ and (○)^2^, and the second layer applies the identity and the exponential function to these powers, (○) and exp(○). The strain energy function *ψ*(***F***) is a sum of the resulting eight terms. Its derivative defines the Piola stress, *∂ψ*(***F***)*/ ≤****F***, whose components, *P*_11_ or *P*_22_, enter the loss function to minimize the error with respect to the biaxial tension data.

Figure 4 shows the *loading* curves for the four plant-based and four animal deli meats directly compared to each other. Percent displacement was prescribed using the CellScale software as 10% in equibiax, for each strip direction, x-stretch in off-x, and y-stretch in off-y for all meats, except the prosciutto meats which were prescribed 25%. For y-str<u>etc</u>h in off ≤ <u>-x an</u>d x-stretch in off-y, the prescribed stretch was 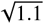 and 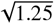 converted to 4.88% and 11.8% relative displacement. As shown in the plots, the instrument did not always reach the prescribed percent displacements when running the tests. The mean is colored in blue for plant-based and red for animal, with the light shading representing the standard error of the mean across the *n* = 8 samples. Plant turkey, plant ham, and plant deli, are the stiffest of the eight meats. Interestingly, plant deli is much less stiff than plant turkey and plant ham, although all three have similar ingredients. Animal prosciutto is the softest of all the meats. The other three animal meats, animal turkey, animal chicken, and animal ham, show a stress-stretch response intermediate between animal prosciutto and the Tofurky deli meats.

**Figure 3:**
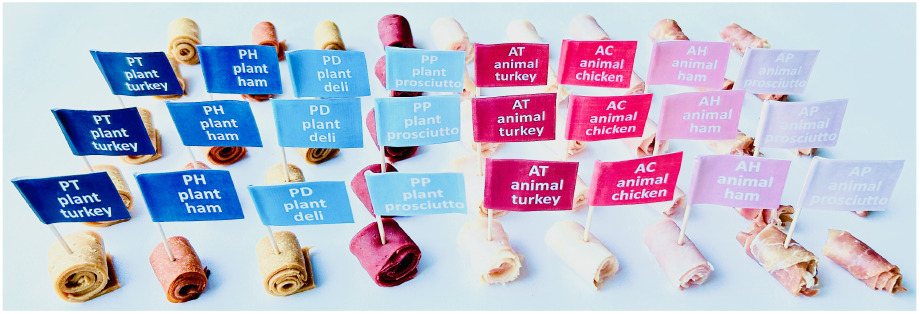
Sensory texture survey samples. Bite-sized samples of the four plant-based deli meats, plant turkey, plant ham, plant deli, and plant prosciutto, and the four animal deli meats, animal turkey, animal chicken, animal ham, and animal prosciutto. Participants rank the sensory features of each product, soft, hard, brittle, chewy, gummy, viscous, springy, sticky, fibrous, fatty, moist, and meaty, on an 5-point Likert scale.

**Figure 4:**
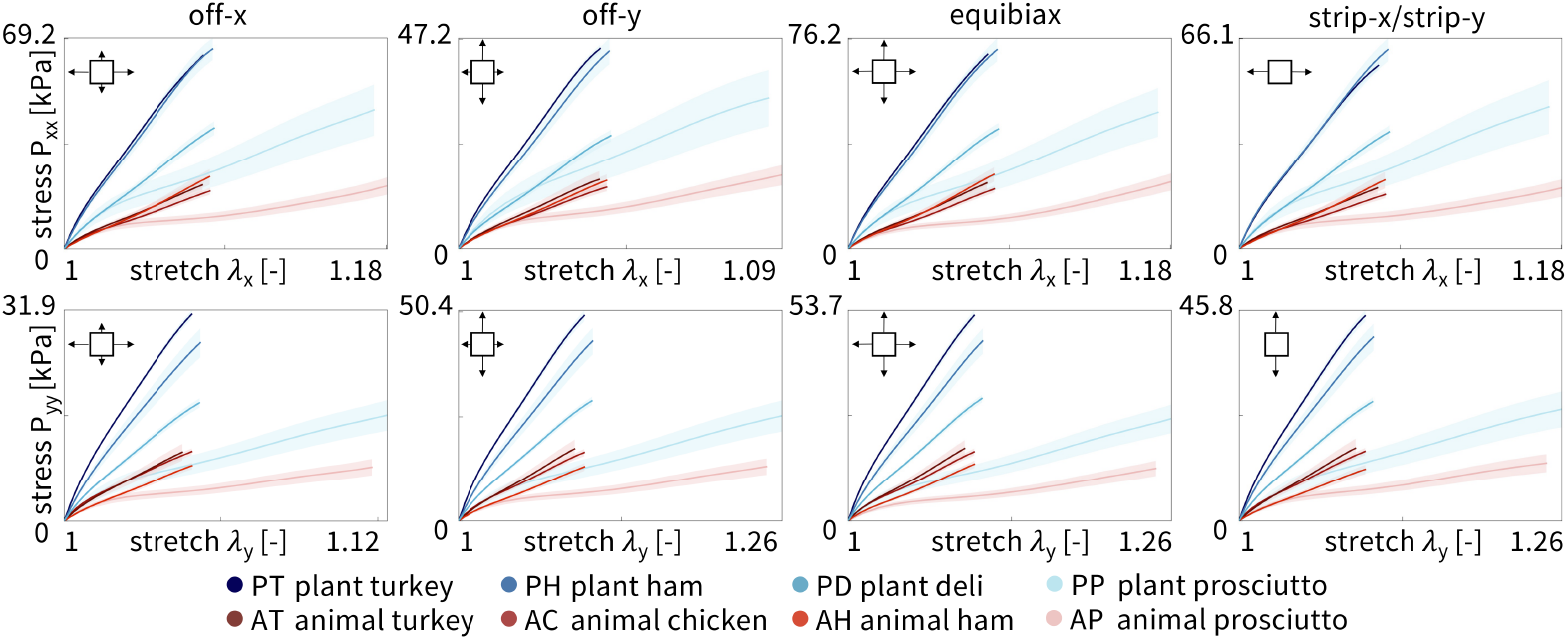
Loading curves of plant-based and animal deli slices for five different biaxial stretch modes: off-x, off-y, equibiax, strip-x, and strip-y. Stress-stretch data are from the third stretch cycle for each mode. The top row shows the stress-stretch curves in the x-direction, and the bottom row shows the stress-stretch curves in the y-direction. Stresses are reported as mean ± standard error of the means of *n* = 8 samples. Plant-based meats are shown in blue and animal meats in red, where curves represent the mean and shading represents the standard error.

Figure 5 shows the *unloading* curves for the four plant-based and four animal deli meats directly compared to each other. Interestingly, all meats exhibited some hysteresis compared to the loading curves, where the unloading curve did not exactly trace back the same curve as the loading response. The stiffest meats in loading, the three Tofurky deli meats, were also the stiffest in unloading. Similarly, the softest meat in loading, animal prosciutto, was also the softest in unloading. Interestingly, plant prosciutto appeared relatively more soft in unloading than in loading compared to animal turkey, animal chicken, and animal ham. We averaged the loading and unloading curves from Figures 4 and 5 to create Table 2.

**Figure 5:**
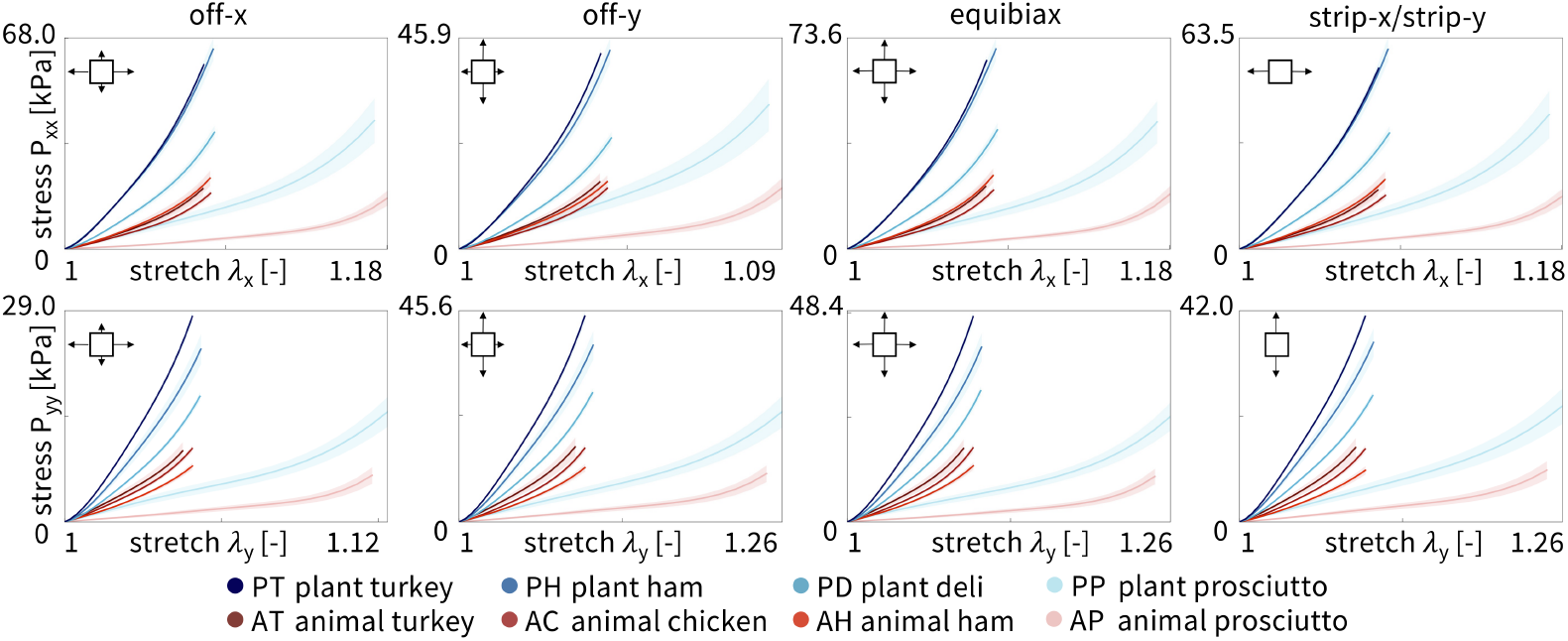
Unloading curves of plant-based and animal deli slices for five different biaxial stretch modes: off-x, off-y, equibiax, strip-x, and strip-y. Data was taken from the third recovery cycle for each mode. The top row shows the stress-stretch curves in the x-direction, and the bottom row shows the stress-stretch curves in the y-direction. Stresses are reported as mean ± standard error of the means of *n* = 8 samples. lant-based meats are shown in blue and animal meats in red, where curves represent the mean and shading represents the standard error.

We fit the parameters of the neo Hooke and Mooney Rivlin models to the average of the loading and unloading stress-stretch curves for each of the *n* = 8 samples of each meat using simple linear regression and report the shear modulus and elastic modulus in Figure 6. The top row shows the shear modulus and the bottom row shows the elastic modulus, E, calculated from the shear modulus *E* = 2*µ*(1 + *v*). Assuming an incompressible material, *v* = 0.5, this simplifies to *E* = 3*µ*. The plots are arranged from highest to lowest value and are the same order for the neo Hooke and Mooney Rivlin models. Darker colors correspond to stiffer meats. Notably, the three tofurky-brand meats are much stiffer than the animal meats. Plant prosciutto has a comparable stiffness to animal turkey, animal chicken, and animal ham, but is twice as stiff as animal prosciutto, the softest meat.

**Figure 6:**
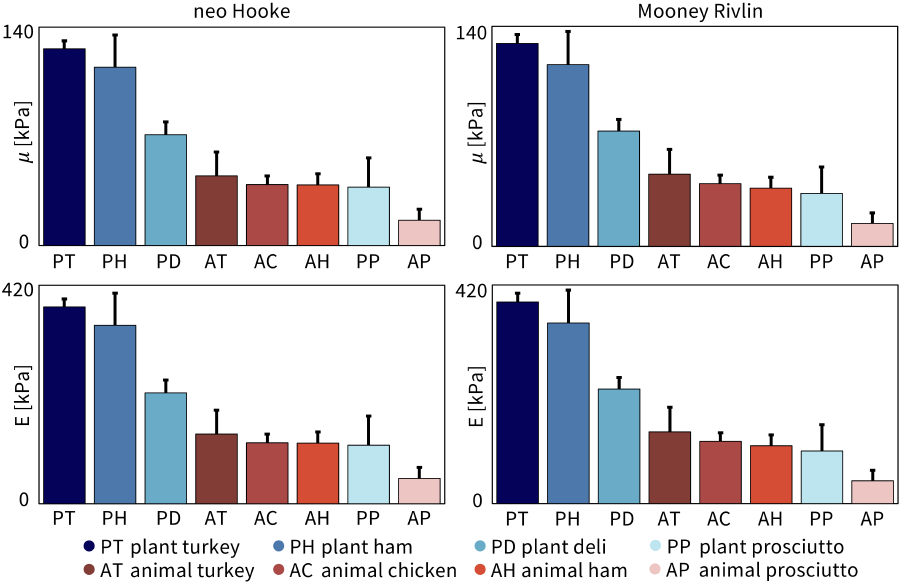
Shear modulus and elastic modulus for all eight deli meats from neo Hooke and Mooney Rivlin models. The top row shows the shear modulus, *µ*, fit using linear regression for the neo Hooke model, left, and Mooney Rivlin model, right. The bottom row shows the elastic modulus, *E*. Parameters are plotted as mean ± standard deviation and arranged in order from highest to lowest, left to right.

Using the data in Table 2 and the constitutive neural network in Figure 2 with L1 regularization, we automatically discover the best two-term models for each of the eight deli meats, shown in Figure 7. The experimental stress is plotted as a function of stretch for each of the five biaxial tension experiments and represented with circles. The colors demonstrate the contributions of the strain energy function, with each color corresponding to a separate model term. The goodness of fit, *R*^2^, is reported for each plot. For strip-x and strip-y, the corresponding hold-y and hold-x are not shown as the stretch is held constant at *λ* = 1. For six out of the eight deli meats, the constitutive neural network discovers the dark red term, (*I*_1_−3). For four of those six meats, (exp(*I*_1_−3) ≤ −1) is the second discovered term. Out of the eight possible model terms, only four, (*I*_1_− 3) and (exp(*I*_1_− 3) ≤ − 1) and (*I*_2_− 3) and (exp(*I*_2_− 3) ≤ − 1), are discovered for any of the meats. Interestingly, the constitutive network discovers exactly the Mooney Rivlin model for animal prosciutto. Only animal ham has neither term from the Mooney Rivlin model but instead takes the exponential of both *I*_1_ and *I*_2_.

**Figure 7:**
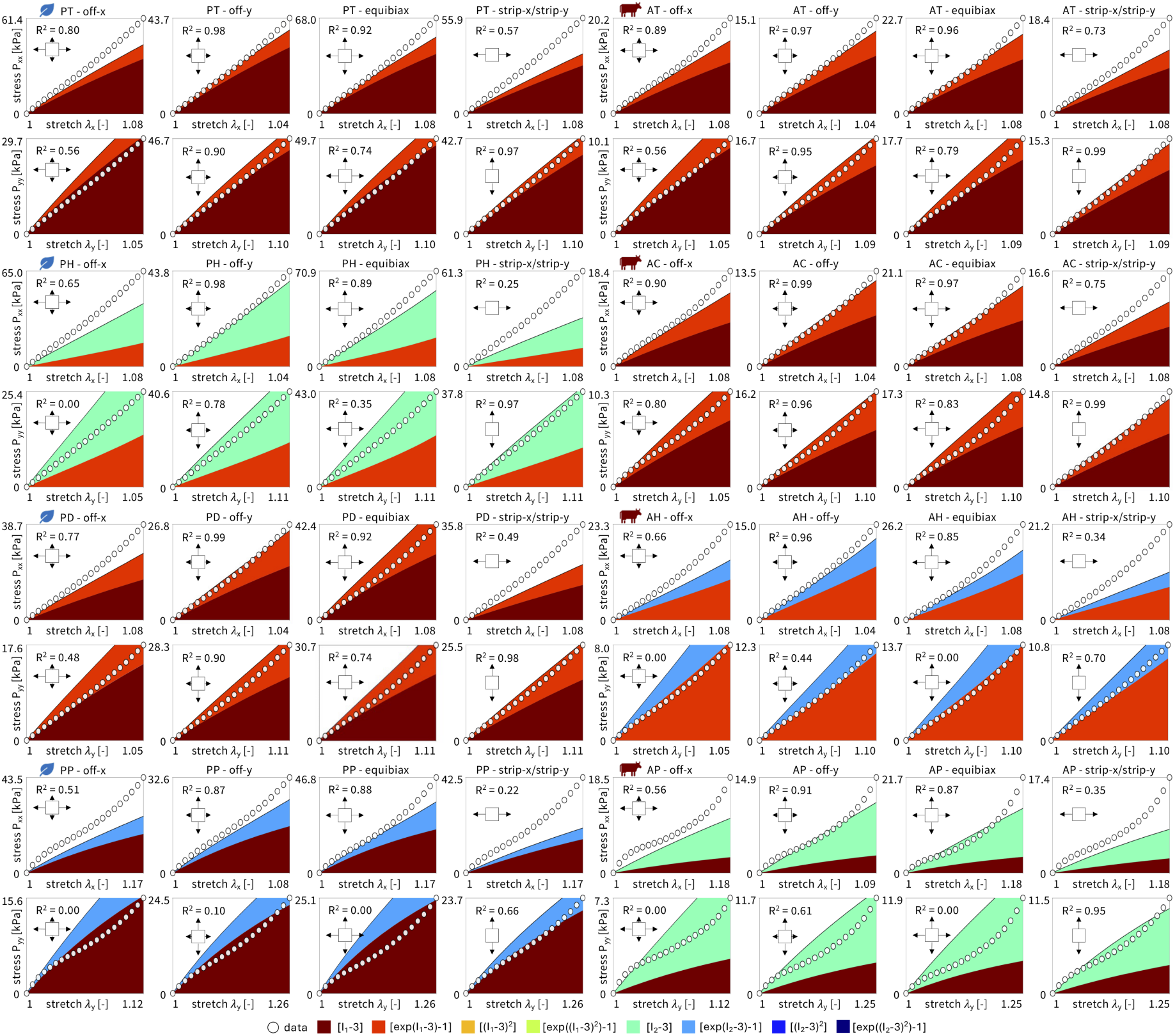
Automated model discovery for all eight deli meats. The constitutive neural network in Figure 2 is trained on all five biaxial loading modes simultaneously for each of the four plant-based and four animal deli meats using the data from Table 1. We apply *L*_1_ regularization to reduce the number of terms to two. The color-coded regions designate the contributions of the eight model terms to the stress function according to Figure 2. The coefficient of determination, *R*^2^, indicates the goodness of fit.

We directly compare how the neo Hooke model, the Mooney Rivlin model, and the newly discovered models fit the experimental biaxial data for each of the eight meats. Figure 8 shows the goodness of fit, *R*^2^, plotted as the mean and standard deviation across the five biaxial loading modes, not including the hold-x and hold-y data from the strip-y and strip-x modes. The meats are arranged in order, from best fitting to worst fitting, with animal chicken the best fitting across all three models with 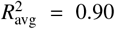 and plant prosciutto the worst fitting with 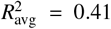. Interestingly, as the fit decreases, the standard deviation error bars generally increase, such that the best fitting meats also have the smallest error bars, and vice versa. Overall, the Mooney Rivlin model provides the best fits across all meats, the newly discovered models are second best, and the neo Hooke model is the worst fitting. However, the differences between the three models are quite marginal with at most a difference of *R*^2^ = 0.1 between the neo Hooke model for plant prosciutto at *R*^2^ = 0.36 and the Mooney Rivlin model at *R*^2^ = 0.46.

**Figure 8:**
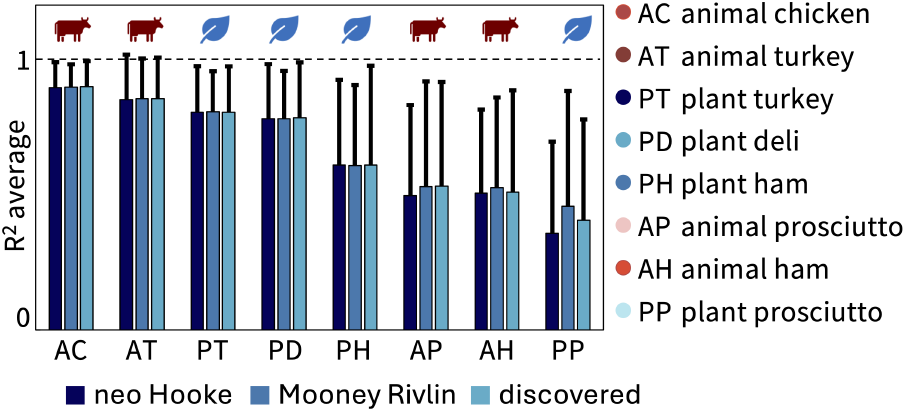
Goodness of fit, *R*^2^, for neo Hooke, Mooney Rivlin, and newly discovered models for all eight deli meats. *R*^2^ indicates the performance of the three models in simultaneously fitting all five biaxial tests. *R*^2^ is plotted as mean ± standard deviation and arranged from best fit, highest *R*^2^, to worst fit, lowest *R*^2^, left to right.

Complementary to the mechanical testing, we surveyed *n* = 18 participants, first, to gain information on how open they are to trying new foods and how attached they are to eating meat, and then, how they perceived certain texture characteristics in deli meat. From our demographic data, 77% of the participants were ages 18-34 and 23% between ages 35-60. The population was 66% white and 56% male. The remaining participants identified as Asian, Hispanic or Latino, American Indian or Alaska Native, Black or African American, female, or nonbinary/gender non-conforming; the exact breakdowns are withheld to protect participant confidentially given the small sample size. Figure 9 shows the results of the Food Neophobia Survey [25] and Meat Attachment Question [8], both of which are validated, pre-existing surveys. The Food Neophobia Survey ranges from 10, *neophilic*, very open to trying new foods, to 70, *neophobic*, not open to trying new foods. The boxplot in dark blue shows the minimum, median, maximum, and first and third quartiles for our participants, while the light blue demonstrates the range of maximum and minimum possible survey results. Our participants were generally quite open to trying new foods, with a median score of 23.5 and a mean score of 24.3. Only one participant skewed more neophobic than neophilic with a score of 42. The Meat Attachment Questionnaire ranges from 16, *not attached* to meat, to 80, *very attached* to eating meat. The boxplot in dark orange shows the minimum, median, maximum, and first and third quartiles for our participants, while the light orange demonstrates the range of maximum and minimum possible survey results. Our participants were slightly more attached to eating meat than not, with a median score of 52 and a mean score of 48.7. Unlike for the Food Neophobia Survey, participants ranged across the entire spectrum of meat attachment, ranging from one participant with the minimum score of 16 to another with a score of 73, nearly the maximum.

**Figure 9:**
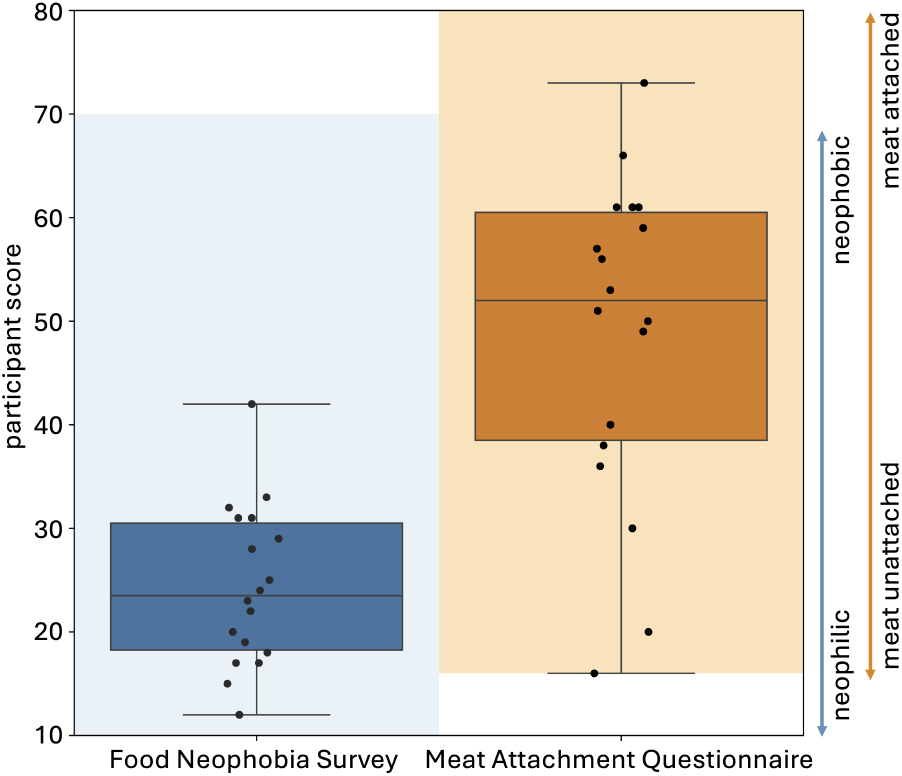
Food Neophobia Survey and Meat Attachment Questionnaire. The Food Neophobia Survey uses a 7-point Likert scale with 10 questions; the light blue region ranges from 10 for neophilic, open to trying new foods to 70 for neophobic, not open to trying new foods [25]. The Meat Attachment Questionnaire uses a 5-point Likert scale with 16 questions; the light orange region ranges from 16 for not attached to eating meat to 80 for very attached to eating meat [8]. The box-and-whisker plots of the minimum, first quartile, median, third quartile, and maximum participant scores are plotted in dark blue and dark orange. The black dots show individual scores.

Figure 10 shows the results of the Sensory Texture Survey, which asks participants to rank each meat from a scale of 1, strongly disagree, to 5, strongly agree, for a series of twelve texture characteristics. These characteristics, *soft, hard, brittle, chewy, gummy, viscous, springy, sticky, fibrous, fatty, moist*, and *meaty* were selected from traditional texture classifications [22, 32] and used in our prior work on three-dimensional minced meat products, primarily hotdogs and sausages [29]. Plots are arranged from most agreement to least agreement. We use a one-way ANOVA to determine if variations between meats for each texture characteristic are significant at *p <* 0.05. Only viscous, springy, and sticky do not have significant variations. Interestingly, three of the twelve characteristics, fibrous, moist, and meaty, split the animal and plant-based deli meats exactly, with all four animal meats ranking higher than all four plant-based meats. Interestingly, plant prosciutto is consistently ranked as comparable to the animal meats, especially for the features soft, fatty, and brittle, where it is the only plant-based meat ranked between the animal meats.

**Figure 10:**
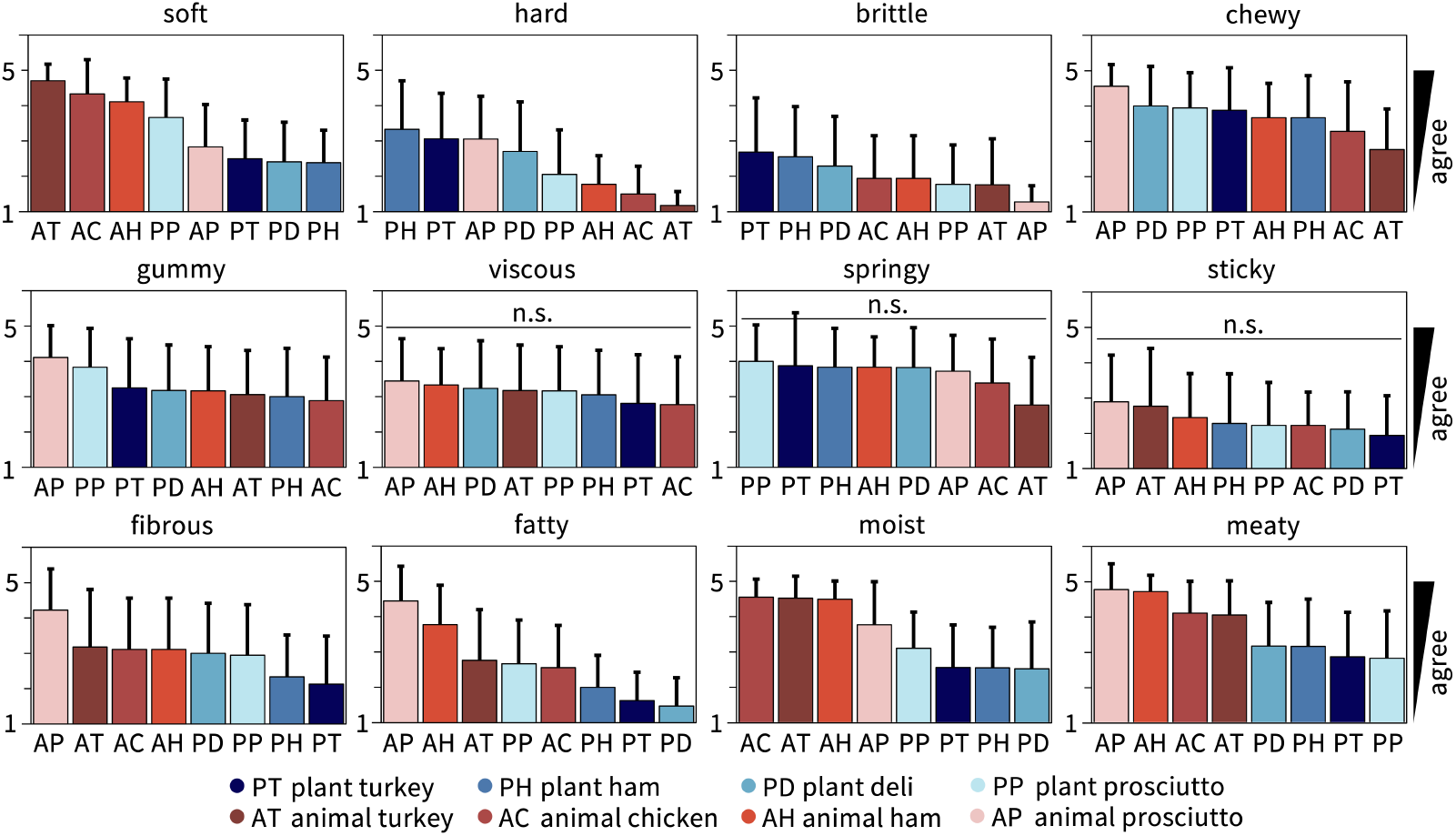
Sensory texture survey results for all eight deli meats. Participants ate a bite-size sample of each deli meat and ranked each texture feature on a 5-point Likert scale ranging from 1, strongly disagree, to 5, strongly agree. Each survey question asked “this food is [texture feature],” with the features soft, hard, brittle, chewy, gummy, viscous, springy, sticky, fibrous, fatty, moist, and meaty adopted from traditional texture classifications [22, 32, 29]. Within each plot, meat products are sorted from highest to lowest agreement, from left to right, with the mean ± standard deviation from *n* = 18 participants plotted. Plant-based meats are colored in blue, animal meats in red. A one-way ANOVA with *p <* 0.05 is used to identify statistically significant variations between meats; n.s. denotes no significant variations.

We created a Spearman’s correlation matrix consisting of the stiffness and the nine significant texture features and found only three significant pairings. First, stiffness is positively correlated with brittleness with *ρ* = 0.857 and *p* = 0.011. Second, stiffness is inversely correlated with fattiness with *ρ* = 0.810 and *p* = 0.022. Lastly, softness is positively correlated with moistness with *ρ* = 0.929 and *p* = 0.002. This contrasts with our prior work with three-dimensional minced meat products, where we found that experimentally measured stiffness was inversely correlated with the perception of softness [29].

## 4. Discussion

We analyzed four plant-based and four animal deli meats using biaxial extension experiments and sensory texture surveys. By comparing mechanical tests with our sensory perception of texture, we can discover how plant-based meats need to improve to better mimic animal meat.

### We discovered the first constitutive model for deli meats

Deli meats are under-researched relative to three-dimensional meat products [1, 16]. As such, no prior constitutive models have been proposed to describe the material behavior of either animal or plant-based deli meats. Here we identified the parameters for the one-term neo Hooke and the two-term Mooney Rivlin models, and used a constitutive neural network to automatically discover the best-fit two-term models out of a library of eight possible terms and 28 possible two-term models. The newly discovered two-term models and the classical Mooney Rivlin model with linear first- and second-invariant terms display nearly identical average fits across the eight deli meats, as we conclude from Figure 8. The classical neo Hooke model with a single linear first-invariant term shows slightly worse fits, although the biggest difference in *R*^2^ was at most 0.1. While four of the meats had an average *R*^2^ above 0.75, the other four meats showed significant variation between the different loading modes, and the one- and two-term models were only able to fit some, but not all of the loading modes at once. Further testing is needed–especially at larger stretches and at different stretch rates–to confirm whether the models and parameters are indeed the best possible models, or more complex models, for example of viscoelastic nature [11, 33], are needed to further reduce the *R*^2^ values. Importantly, our study not only characterizes the mechanics of deli meat in terms of a *single parameter*, for example the stiffness or shear modulus, but also it discovers *fully three-dimensional models and parameters* that enable physics-based simulations and predictions of each meat’s material behavior. Simulations allow for virtual experiments with different ingredients and formulations and can even virtually mimic the act of chewing, drastically speeding up the design of new products and decreasing the cost and time to market [24, 27].

### Physical stiffness is correlated with sensory brittleness

No study to date has compared the physical properties and sensory texture of plant-based and animal deli meats. In a direct side-by-side comparison, we observe that our participants were able to correlate the *physical stiffness* with the *sensory brittleness*, with Spearman’s correlation coefficient of *ρ* = 0.857 and *p* = 0.011, but not with the sensory softness or hardness. In a previous study of three-dimensional minced meat products, we found that the physical stiffness, measured across tension, compression, and shear tests, was significantly correlated with hardness and inversely correlated with softness [29]. It seems intuitive that hardness and softness are more difficult to perceive for thin deli meat slices than for three-dimensional cubical samples of meat. For instance, although the animal prosciutto slices are physically less than half as stiff as the other animal deli slices, participants ranked their sensory perception as harder and less soft. These discrepancies may be explained, at least in part, because we worked with a small untrained consumer panel, rather than with a trained panel of sensory experts. Yet, trained sensory panels, while valuable, are still subjective, variable, and influenced by personal preferences, training, and environmental factors [16]. Future studies with larger sample sizes are needed to determine which textural characteristics represent truly distinguishable features of deli meat. Taken together, our observations emphasize the need for mechanical testing as an objective, quantitative, and reproducible alternative to eliminate the inherent biases of sensory panel surveys. Mechanical stiffness testing ensures consistency across samples–independent of human perception–and provides precise, high-resolution data for accurate product development and product optimization.

### Current plant-based deli meats fail to mimic the physical and sensory signature of animal deli meats

When designing new plant-based meats, mimicking the physical and sensory texture of animal meat is critical for consumer acceptance, since texture strongly influences our perception of taste and overall eating experience. Strikingly, the plant-based products, turkey, ham, deli, and prosciutto, with physical stiffnesses of 378 ± 15 kPa, 343 ± 62 kPa, 213 ± 25 kPa, and 113 ± 56 kPa, were *more than twice as stiff* as their animal counterparts, turkey, chicken, ham, and prosciutto, with 134 ± 46 kPa, 117 ± 17 kPa, 117 ± 21 kPa, and 49 ± 21 kPa, as we conclude from Table 1 and Figure 6. This mismatch is reflected in the sensory survey, in which the participants perceived all four plant-based products as *less fibrous, less moist*, and *less meaty* than the four animal products, as we conclude from Figure 10. Of all plant-based products, plant-based prosciutto with a physical stiffness of 113 ± 56 kPa comes closest to the four animal products. Its sensory perception of softness, hardness, brittleness, and fattiness consistently ranks within the feature rankings for animal meats. Interestingly, although the other three plant-based products, plant-based ham, plant-based turkey, and plant-based deli, share the same first three ingredients–water, wheat gluten, and organic tofu–their stiffnesses vary by almost a factor two. Of these three, plant-based deli has the closest physical stiffness to the animal meats. Interestingly, it was also ranked the most fibrous and the most meaty of the plant-based meats. Our study confirms the common belief that current plant-based deli products struggle to meet the physical stiffness and sensory perception of animal deli meat. More research and development is needed to fully explore the parameter space of ingredients, formulations, and processing methods to accurately replicate the texture of animal products [30].

## 5. Conclusion

Biaxial testing with multiple different stretch ratios reveals the complex material behavior of four plant-based and four animal deli meats. Here we identified material models and parameters that best describe the relation between stress and stretch for all eight products. We observed that the plant-based turkey, ham, deli, and prosciutto, were more than twice as stiff as their animal counterparts, turkey, chicken, ham, and prosciutto. Sensory texture surveys reveal how people perceive these differences in physical stiffness. Our survey participants ranked all four plant-based products as less fibrous, less moist, and less meaty than the four animal products. Our study reveals that the physical stiffness of deli meat is correlated to the sensory perception of brittleness, but not to hardness or softness. Of all plant-based products, plant prosciutto comes closest to the animal products, both in physical stiffness and in sensory softness, hardness, brittleness, and fattiness. Taken together, current plant-based deli slices struggle to replicate the complete physical and sensory signature of animal deli meats. We anticipate that, by integrating mechanical testing, sensory panels, and artificial intelligence, we can identify the best combinations of ingredients, formulations, and processing methods to design plant-based meat alternatives that are delicious, nutritious, and more environmentally friendly than traditional animal meat.

## Data Availability

Data, constitutive neural network code, and linear regression analysis script are freely available at https://github.com/LivingMatterLab/CANN.

## Acknowledgments

This work was supported by seed funding from the Stanford Plant-Based Diet Initiative and from Food System Innovations, by the NSF Graduate Research Fellowship to Skyler R. St. Pierre and Ethan C. Darwin, by the Stanford DARE Fellowship to Skyler R. St. Pierre, by the NSF CMMI Award 2320933 Automated Model Discovery for Soft Matter, and by the ERC Advanced Grant 101141626 DISCOVER to Ellen Kuhl.

